# Two distinct classes of co-chaperones compete for the EEVD motif in heat shock protein 70 (Hsp70) to tune its activity

**DOI:** 10.1101/2021.10.18.464838

**Authors:** Oleta T. Johnson, Cory M. Nadel, Emma C. Carroll, Taylor Arhar, Jason E. Gestwicki

## Abstract

Chaperones of the heat shock protein 70 (Hsp70) family engage in protein-protein interactions (PPIs) with many co-chaperones. One hotspot for co-chaperone binding is the EEVD motif that is found at the extreme C-terminus of cytoplasmic Hsp70s. This motif is known to bind tetratricopeptide repeat (TPR) domain co-chaperones, such as the E3 ubiquitin ligase CHIP, and Class B J-domain proteins (JDPs), such as DnaJB4. Although complexes between Hsp70-CHIP and Hsp70-DnaJB4 are both important for chaperone functions, the molecular determinants that dictate the competition between these co-chaperones are not clear. Using a collection of EEVD-derived peptides, we find that DnaJB4 binds to the IEEVD motif of Hsp70s, but not the related MEEVD motif of cytoplasmic Hsp90s. Then, we explored which residues are critical for binding to CHIP and DnaJB4, revealing that they rely on some shared features of the IEEVD motif, such as the C-terminal carboxylate. However, they also had unique preferences, especially at the isoleucine position. Finally, we observed a functionally important role for competition between CHIP and DnaJB4 *in vitro*, as DnaJB4 can limit the ubiquitination activity of the Hsp70-CHIP complex, while CHIP suppresses the chaperone activities of Hsp70-DnaJB4. Together, these results suggest that the EEVD motif has evolved to support diverse PPIs, such that competition between co-chaperones could help guide whether Hsp70-bound proteins are folded or degraded.

## Introduction

Members of the heat shock protein 70 (Hsp70) family of molecular chaperones play a critical role in maintaining protein homeostasis (aka proteostasis). Hsp70s are composed of a nucleotide binding domain (NBD), a substrate binding domain (SBD), and a C-terminal unstructured region terminating in an EEVD motif (Fig 1A)^1,2^. A major structural feature of Hsp70s is that ATPase activity in the NBD causes a conformational change that allosterically regulates the affinity of the SBD for “client” proteins^3,4^. This general binding mechanism allows Hsp70s to recognize a wide array of clients and then to function in diverse processes such as protein folding, translocation, complex formation, and degradation^5,6^.

**Figure 1:**
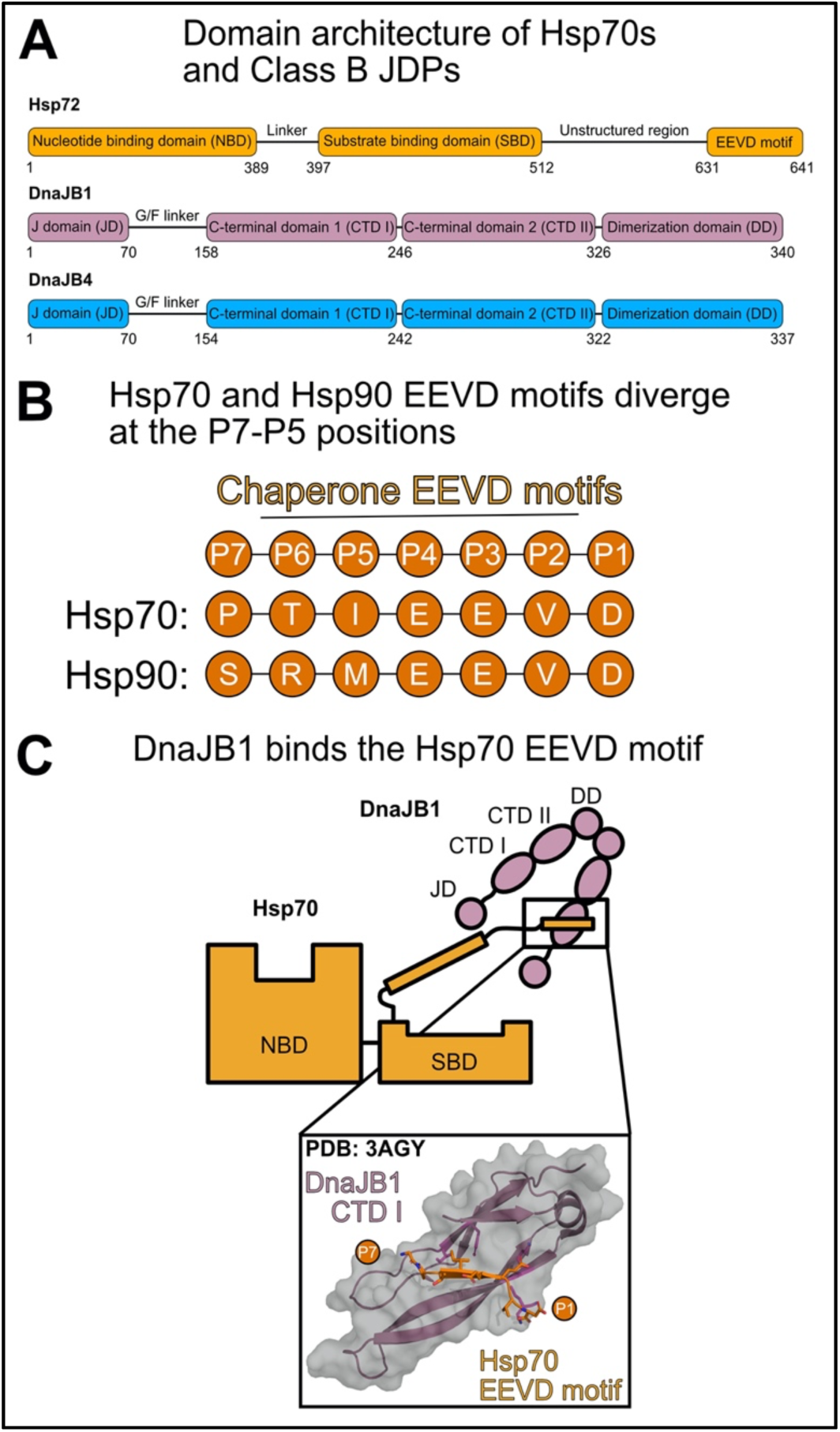
Summary of the interactions between class B JDPs and the C-terminal EEVD motif of Hsp70. (A) Domain architecture of Hsp72 and the canonical, Class B JDPs: DnaJB1 and DnaJB4. (B) Position nomenclature for the EEVD motifs of Hsp70 (Hsp72/HSPA1A) and Hsp90 (Hsp90α*/*HSP90AA1). In this nomenclature, the C-terminal aspartate is termed P1, such that the sequences of Hsp70s and Hsp90s begin to diverge in the P5 through P7 positions. (C) Cartoon and crystal structure representations of the interaction between Hsp70’s EEVD motif (orange) and DnaJB1’s CTD I (purple; PDB 3AGY^37^). Not shown in the cartoon schematic is the important interaction of the JD with Hsp70’s NBD and SBD. Also, binding to only one CTD I in the dimer is shown for simplicity, but both are likely competent for this interaction.

However, Hsp70s rarely work alone. Rather, the diversity of Hsp70’s functions is imparted by co-chaperones, such as J-domain proteins (JDPs)^7,8^, nucleotide exchange factors (NEFs)^9^, and tetratricopeptide repeat (TPR) domain proteins^10^. Some of these co-chaperones, such as JDPs and NEFs, bind Hsp70s and stimulate cycles of nucleotide hydrolysis to regulate client binding^11–14^. In addition, co-chaperones also act as adaptors, connecting Hsp70s and their clients to other cellular effector functions. For example, some NEFs and TPR proteins link Hsp70’s clients to protein degradation pathways^15,16-17^. Thus, collaboration between Hsp70s and their co-chaperones, mediated by a series of direct protein-protein interactions (PPIs), is critical for establishing the functional diversity of the chaperone^19,20^. It seems that a key feature of this system, therefore, is that there are limited surfaces on Hsp70s for co-chaperones to bind, such that the co-chaperones must compete for shared sites. In turn, the recruitment of a specific co-chaperone over others will, in part, dictate what happens to the client. Accordingly, it is important to understand how Hsp70 binds its different co-chaperones and what molecular features drive those decisions.

A major “hotspot” of co-chaperone binding is the EEVD motif that is present at the extreme C-terminus of cytoplasmic Hsp70s^21,22^. Interestingly, cytoplasmic Hsp90s, despite their dramatically different overall structure, also contain a conserved C-terminal EEVD motif^23^. However, the Hsp70 and Hsp90 motifs are not identical, as they differ at positions N-terminal to the four residue, EEVD sequence (Fig 1B). Here, we will use nomenclature in which the C-terminal aspartate is termed P1, while P2 is the valine, *etc*. In this systemization, the cytoplasmic Hsp70s have an isoleucine at P5 (IEEVD), while the cytoplasmic Hsp90s have a methionine (MEEVD).

The interactions between the EEVD motifs and TPR co-chaperones have been extensively characterized^24–27^. Co-crystal structures of EEVD peptides bound to representative TPR domains have shown that a striking feature of this interaction is a “carboxylate clamp” in the TPR domain, involving lysine and arginine side chains, that coordinates the P1 aspartic acid and the EEVD’s carboxy-terminus^28,29^. Biochemical studies have also shown that the identity of the P5 residue is also important, such that some co-chaperones have a strong preference for Hsp90’s MEEVD over Hsp70’s IEEVD^30–35^. This is an important finding because it shows that relatively small changes in the sequence (*e*.*g*. isoleucine *vs*. methionine at P5) can have dramatic consequences for TPR binding preferences. This idea has been taken even further in studies that have used systematic mutations in Hsp70’s IEEVD motif to reveal the deeper structure-activity relationships (SAR) for binding to certain TPR proteins, such as HOP and CHIP^34,36^. For example, these SAR studies have revealed that the identity of the P2 residue is important for CHIP binding, but that both conserved P3 and P4 glutamates are dispensable^36^. Together, these structural and biochemical studies have started to reveal how TPR co-chaperones “read” chemical information in the EEVD motifs to generate functionally distinct complexes.

The EEVD motif is also the site for binding another subclass of co-chaperones, the Class B JDPs^37–39^. JDPs are named for their conserved J-domain (JD)^40–43^, which binds Hsp70s near the interdomain linker between the NBD and SBD^44^. This interaction requires an invariant HPD sequence within the JD and it is responsible for the stimulation of Hsp70’s ATPase activity^7,8,11,45^. Outside of the conserved JD, however, the members of the JDP family vary in their structure and domains. Broadly, these differences have led to the JDPs being placed into 3 structural categories (Class A, B and C). Here, we focus on the Class Bs because they are the only ones shown to bind the EEVD motif. The Class B JDPs are typified by a glycine-phenylalanine rich linker (G/F), two beta-barrel domains termed C-terminal domains 1 and 2 (CTD I/CTD II), and a dimerization domain (DD) (Fig 1A)^8^. Initially, the CTD I and CTD II domains were found to interact with prospective Hsp70 clients^38,46–48^, serving to recognize and deliver them to Hsp70s. However, later work found that CTD I is also the site of interaction with the EEVD motif^46,49^. This interaction was first characterized in the yeast JDP, Sis1, as well as the human JDP, Hdj1/DnaJB1 (Fig 1C, PDB 3AGY)^37–39,50,51^. That work showed that, although the PPI between CTD I and the EEVD motif is weak (Kd ≅10-20 µM)^38,52^, it is functionally important for coordinating with Hsp70’s functions. For example, when the EEVD interaction is impaired, the ability of the Hsp70 system to fold clients is inhibited^21,38,39,53,54^. The reasons for this dependence have been elucidated by NMR studies, which have shown that the EEVD interaction at CTD I allosterically relieves autoinhibition of the JD by the G/F linker, allowing engagement of the JD with Hsp70 and promoting ATPase activity, client refolding, and disaggregation of amyloids^39,55^.

Compared to the binding of TPR domains to the EEVD motif, less is known about the molecular determinants of the JDP-EEVD interaction. Specifically, while structural studies have identified residues that are critical to binding^37–39,50,53,54^, a detailed SAR has not yet been described. We reasoned that this gap in knowledge limits our understanding of how TPR and JDP co-chaperones might compete for EEVD motifs to tune chaperone functions. Here, we characterize the determinants of the JDP-EEVD interaction using the representative Class B JDP, DnaJB4. We chose to focus on DnaJB4 because, while both DnaJB4 and DnaJB1 are known to collaborate with Hsp70 in functional assays^56^ and structural studies have been performed on DnaJB1, comparatively little is known about the binding of EEVD motifs to DnaJB4. Using fluorescence polarization (FP) and differential scanning fluorimetry (DSF), we found that DnaJB4 binds selectively to the Hsp70 IEEVD motif, but not the Hsp90 MEEVD sequence. Using truncations and mutations, we also found that DnaJB4 recognizes the carboxy-terminus and has strong preferences for the P5 residue. Based on this knowledge, we developed an inactivating point mutation in Hsp70’s EEVD and used it to probe the functional importance of the interaction, showing that it is critical for both ATPase and client refolding activities. We also confirmed that DnaJB4 can interfere with the function of the Hsp70-CHIP complex in ubiquitination assays and that, conversely, CHIP can partially disrupt the chaperone functions of the Hsp70-DnaJB4 complex. Finally, we found that pseudo-phosphorylation of the EEVD motif selectively interrupts binding to CHIP, but leaves the interaction with DnaJB4 intact. Together, these studies suggest that competition between distinct classes of co-chaperones can tune the function of Hsp70 complexes.

## Results

### DnaJB4 binds Hsp70’s C-terminal EEVD motif

Structural studies have shown how the EEVD motif binds to CTD I of DnaJB1 (see Fig 1C). Given the conservation between the CTD I domains of DnaJB1 and DnaJB4 (Figure 2A), we hypothesized that DnaJB4 may bind the EEVD motif in a similar way. To explore this possibility, we first aligned a predicted structure of DnaJB4’s CTD I generated by AlphaFold v2.0^57^ to the crystal structure of DnaJB1 bound to the C-terminal peptide from Hsp72/HSPA1A (PTIEEVD-COOH) (PDB 3AGY^37^). An overlay of the two domains showed that key interactions with were predicted to be conserved (Fig 2A). Notably, cationic residues that coordinate the P3 glutamate sidechain (DnaJB1 Lys181, DnaJB4 Lys177) and the EEVD carboxy-terminus (DnaJB1 Lys182, DnaJB4 Arg178) were structurally conserved in both CTD I domains. We also identified a series of conserved hydrophobic contacts surrounding the sidechain of the P5 isoleucine (DnaJB1 Ile235/Phe237 or DnaJB4 Ile231/Phe233). Other sidechains in the EEVD, such as P1, P2 and P6, were solvent exposed and perhaps not as likely to make direct interactions.

**Figure 2:**
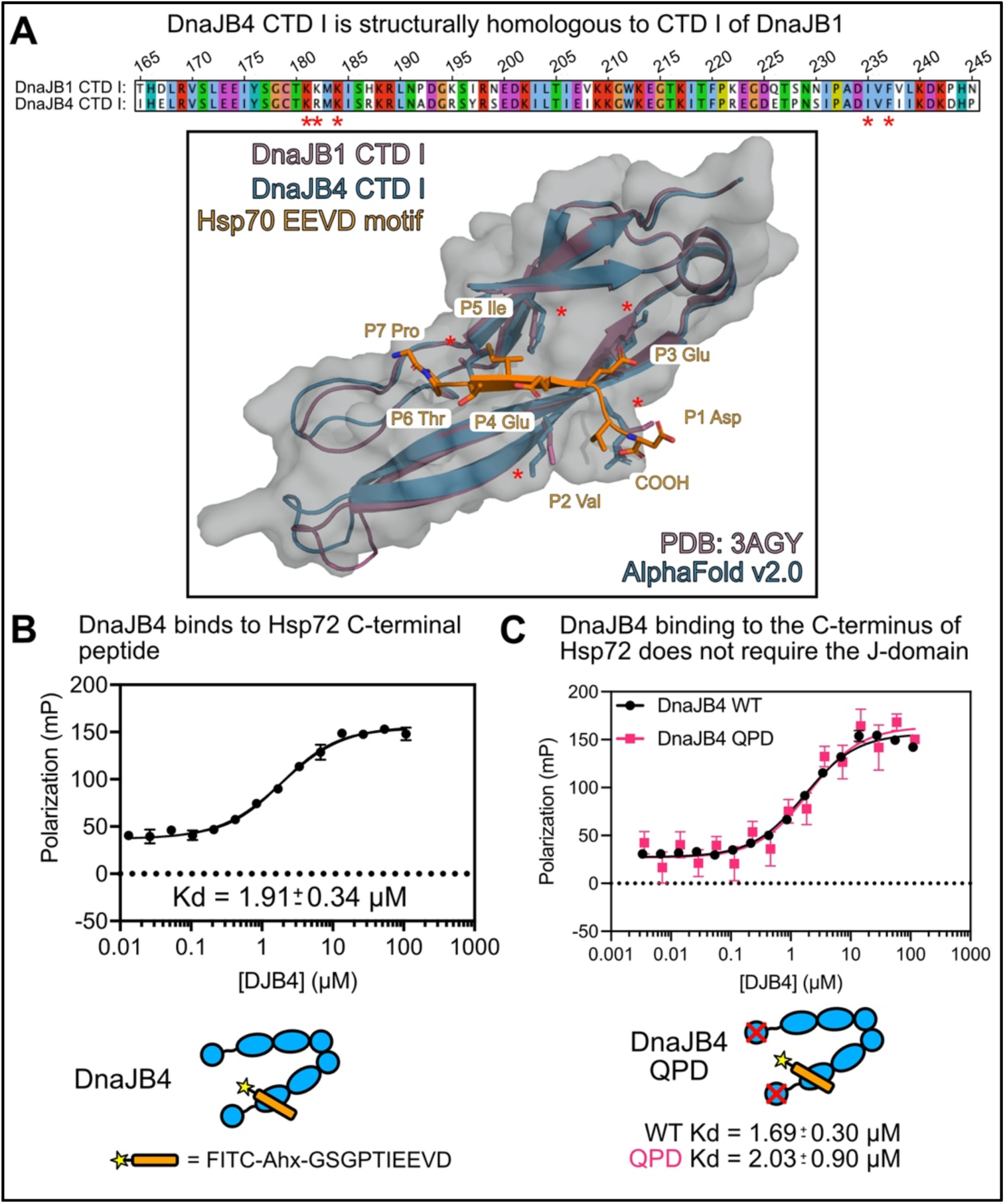
DnaJB4 binds Hsp70’s C-terminal EEVD motif. (A) Sequence alignment and structural overlay of the CTD I domains of DnaJB1 and DnaJB4. Critical residues for binding to Hsp70’s EEVD motif are highlighted by red asterisks. Crystal structure of DnaJB1 CTD I in complex with Hsp70 EEVD motif (PDB 3AGY^37^, DnaJB1 = purple, Hsp70 EEVD = orange) was aligned to the predicted structure of DnaJB4 CTD I (AlphaFold v2.0^57^, Blue) in PyMol v2.x (Schrodinger). (B) Saturation binding of Hsp72 tracer to recombinant, human DnaJB4, as measured by FP. The tracer sequence is included (bottom). The results are the average of four replicates and error bars represent standard deviation (SD). Some error bars are smaller than the symbol. The Kd is shown as mean with a 95% confidence interval (CI). (C) Saturation binding of Hsp72 tracer binding to WT DnaJB4 or the QPD mutant. The results are the average of four replicates and error bars represent SD. Some error bars are smaller than the symbol. Kd values are shown as the mean with a 95% CI.

To test the prediction that the EEVD motif would bind DnaJB4, we created a fluorescently labelled peptide derived from the last 10 residues of Hsp72 (*FITC-Ahx-GSGPTIEEVD*) and measured binding to recombinant, full length DnaJB4 by fluorescence polarization (FP). In this platform, DnaJB4 bound with a Kd of 1.91 ± 0.34 µM (Fig 2B). To reveal any contribution of the J-domain to this interaction, we introduced the well-known QPD mutation^58^. As expected, this mutation had no effect on binding to the EEVD tracer, as both WT and the QPD DnaJB4 mutant bound with comparable affinities (WT Kd = 1.69 ± 0.03 µM; QPD Kd = 2.03 ± 0.90 µM) (Fig 2C).

### DnaJB4 binds Hsp70’s IEEVD motif, but not the closely related MEEVD from Hsp90

To probe whether DnaJB4 could also bind the Hsp90 MEEVD motif, we generated unlabeled, *N*-acetylated 10mer peptides corresponding to the C-termini of the most abundant cytoplasmic chaperones from the Hsp70 (Hsc70/HSPA8, Hsp72/HSPA1A) and Hsp90 (Hsp90α, and Hsp90β) classes (Fig 3A). Then, binding to DnaJB4 was measured using differential scanning fluorimetry (DSF). In the absence of peptide, DnaJB4 showed an apparent melting temperature (Tm_app_) of 57 ± 0.06 °C (Fig 3B). Treatment with peptides from either Hsp72 or Hsc70 caused a positive shift of approximately 1 to 2 °C in the melt curves (Hsp72 Tm_app_ = 58 ± 0.21 °C; Hsc70 Tm_app_ = 59 ± 0.14 °C), suggesting that they bind to DnaJB4. Conversely, neither Hsp90α nor Hsp90β peptides bound to DnaJB4 (Hsp90α Tm_app_ = 57 ± 0.15 °C; Hsp90β Tm_app_ = 57 ± 0.20 °C). To confirm this result, we tested the same peptides as competitors in the FP assay, using the fluorescent Hsp72 peptide as the tracer. In this format, peptides from Hsp72 and Hsc70 again bound to DnaJB4 (% tracer displacement Hsp72 = 40.2 ± 4.6%, Hsc70 = 67.6 ± 4.4%), while Hsp90α and Hsp90β peptides did not (Hsp90α = -7.7 ± 1.5%, Hsp90β -1.3 ± 2.2%) (Fig 3C). To specifically ask whether the P5 Ile/Met contributes to this dramatic difference in binding, we substituted the P5 Ile in the Hsp72 10-mer for Met and then measured binding to DnaJB4 by FP. Indeed, this single mutation was sufficient to weaken binding to DnaJB4 (Fig 3D). Taken together, these data show a direct interaction between DnaJB4 and chaperone C-termini that is selective for the Hsp70 system.

**Figure 3:**
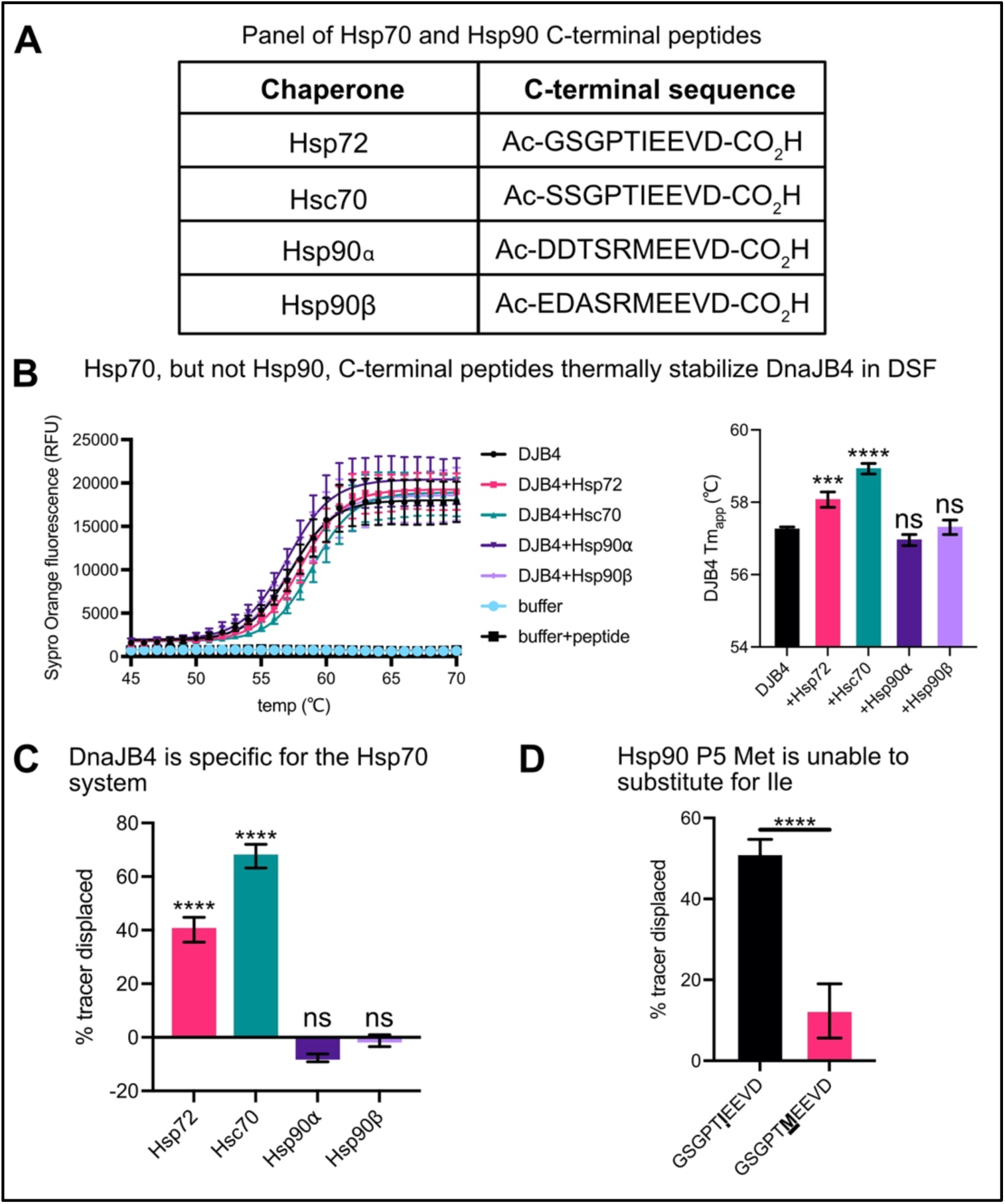
DnaJB4 binds Hsp70’s IEEVD motif, but not the closely related MEEVD from Hsp90. (A) Table listing the sequences of chaperone C-terminal peptides used in this study. (B) DSF melting curves and apparent melting temperatures (Tm_app_) of 5 µM DnaJB4 in the presence of either a DMSO control or various chaperone peptides (100 µM). Temperature-dependent unfolding was monitored by Sypro Orange (SO) fluorescence. The melting curves represent the mean SO fluorescence ± SD (n=4). Buffer alone and buffer + peptide samples were used as negative controls. The calculated DnaJB4 Tm_app_ values are mean ± SD (n=4). Statistics were performed using unpaired student’s t-test (***p<0.001, ****p<0.0001, ns = not significant compared to DMSO control). (C) FP experiment showing displacement of Hsp72 probe from DnaJB4 by various chaperone competitor peptides. Graph shows the mean tracer displacement relative to a DMSO control ± SD (n=4). Statistics were performed using unpaired student’s t-test (***p<0.0001, ns = not significant compared to DMSO control). (D) Competition FP experiment comparing WT Hsp72 to a mutant in which the P5 Ile was replaced by Met. Graph shows the mean tracer displacement relative to DMSO control ± SD (n=4). Statistics were performed using unpaired student’s t-test (****p<0.0001 compared to WT control).

### Residue-level determinants of the JDP-EEVD interaction

After establishing that DnaJB4 interacts with the Hsp70 C-terminus, we wanted to further delineate the molecular determinants of this PPI. The C-terminal carboxylate is known to be obligate for EEVD binding to TPR domain co-chaperones, such as CHIP^36^ ; thus, we first asked whether the carboxylate might also be required for binding to DnaJB4. In FP competition assays, amidation of the carboxylate drastically reduced binding (Fig 4A), suggesting that it is indeed a critical feature. Next, we used truncations to probe how much of the 10-mer sequence was required. Interestingly, removing the P10, P9 or P8 positions improved, rather than inhibited, binding to DnaJB4 (Fig 4B). Truncation of the P7 Pro residue (TIEEVD), however, significantly lowered affinity, while truncation of the P6 Thr residue (IEEVD) further weakened it. These findings are consistent with the predicted interactions in the DnaJB4-bound complex (see Fig 2A), wherein the last 7 amino acids of Hsp70’s C-terminus (PTIEEVD) are necessary to span the entire PPI interface on DnaJB4’s CTD I.

**Figure 4:**
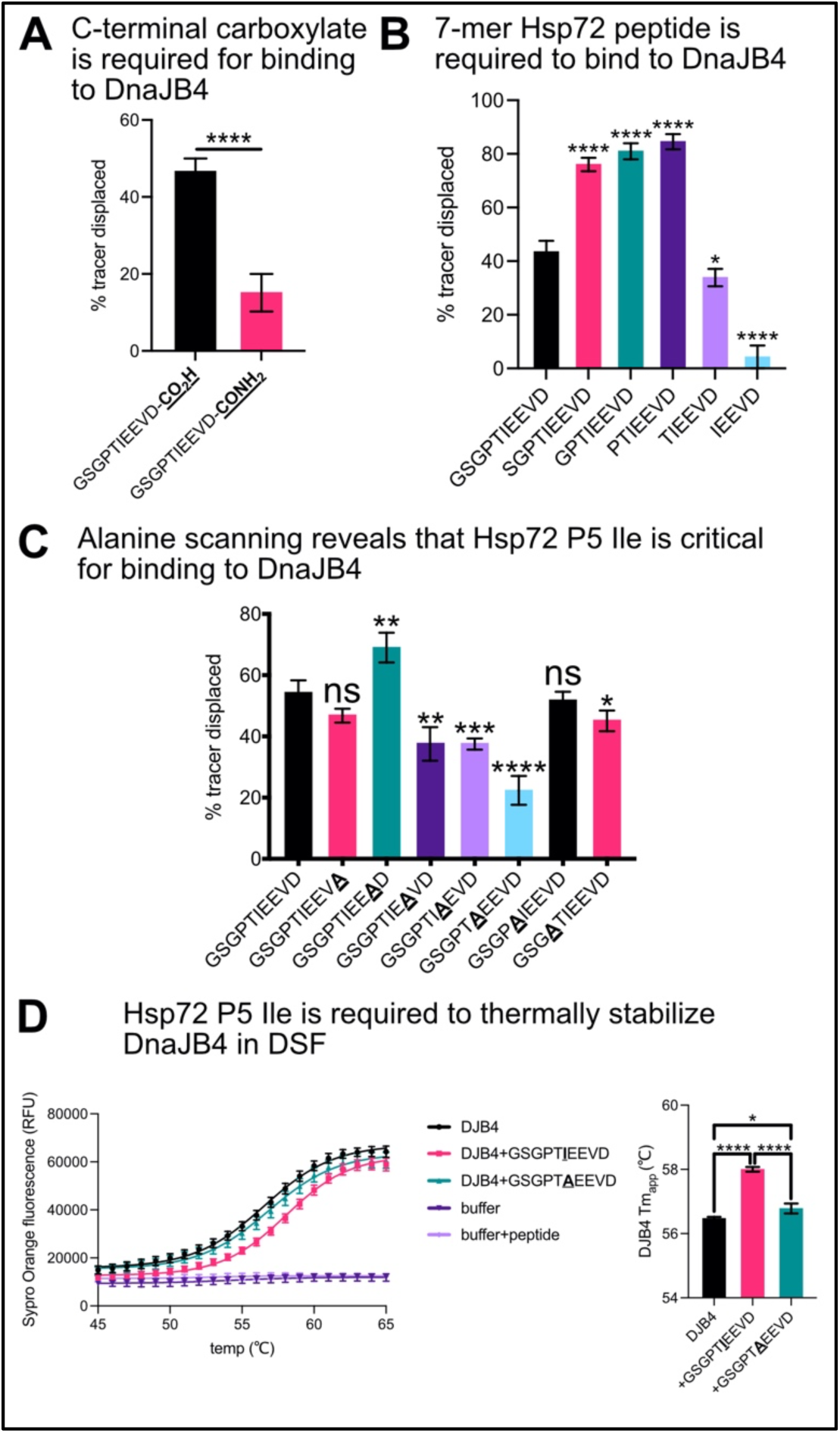
Residue-level determinants of the JDP-EEVD interaction. (A) Competition FP experiment comparing displacement of Hsp72 tracer from DnaJB4 by WT or C-terminally amidated Hsp72 peptides. Graph shows mean tracer displacement relative to DMSO control ± SD (n=4). Statistics were performed using unpaired student’s t-test (****p<0.0001 compared to WT control). (B) Competition FP experiment comparing displacement of Hsp72 tracer from DnaJB4 by N-terminally truncated competitor peptides. Graph shows mean tracer displacement relative to DMSO control ± SD (n=4). Statistics were performed using unpaired student’s t-test (*p<0.05, ****p<0.0001 compared to WT control). (C) Competition FP experiment comparing alanine scanning substitutions across the Hsp72 C-terminal sequence in binding to DnaJB4. Graph shows mean tracer displacement relative to DMSO control ± SD (n=4). Statistics were performed using unpaired student’s t-test (*p<0.05, **p<0.01, ***p<0.001, ****p<0.0001, ns = not significant compared to WT control). (D) DSF experiment confirming the role of the P5 Ile in Hsp72 EEVD motif binding to DnaJB4. Temperature-dependent unfolding was monitored by SO fluorescence at the denoted temperatures (°C). Melt curves are representative of the mean SO fluorescence in RFU ± SD (n=4). Buffer alone and buffer + peptide samples were used as negative controls. DnaJB4 Tm_app_ is represented as mean ± SD (n=4). Statistics were performed using unpaired student’s t-test (*p<0.05, ****p<0.0001).

Having identified that the P1-P7 residues are required for the interaction with DnaJB4, we then performed an alanine scan of the 7-mer sequence to assess the contributions of each residue. The most dramatic effect was found at P5, where an alanine mutation significantly weakened binding (Fig 4C). We confirmed this result using DSF, finding that the single P5 Ile to Ala mutation abrogated thermal stabilization of DnaJB4 (Fig 4D). This result can be rationalized in the predicted structure, where the Hsp72 P5 isoleucine is “caged” by neighboring hydrophobic residues, Ile231 and Phe233, in DnaJB4 (see Fig 2A). The only other positions that were sensitive to alanine mutation were the two glutamates at P3 and P4, with a more modest effect on the P7 proline. Together, these studies suggest that the carboxylate and the P5 Ile are most important for binding to DnaJB4 and that other side chains, especially P3, P4 and P7, make additional contributions to equilibrium binding affinity.

### DnaJB4 accommodates an expanded number of amino acids at P5, compared to CHIP

It is well known that TPR domain co-chaperones recognize the P5 residue and the carboxy-terminus in the EEVD motif^2,25,59^. Because we found that the same positions are also critical for binding DnaJB4 (see Fig 4), we expected that TPR co-chaperones, such as CHIP, would compete for binding (as schematized in Fig 5A). However, it was still not clear whether DnaJB4 and CHIP would have the same sequence requirements at P5. As an initial step in asking this question, we first confirmed that the amidated Hsp72 C-terminus (GSGPTIEEVD-CONH_2_) was unable to bind CHIP in our FP assay (Fig 5B). Then, we used the alanine scan peptides to show that the P1, P2 and P5 positions are indeed most important for binding CHIP (Fig 5C). These conclusions matched well with previous benchmark studies^36^, but it was important for us to repeat them with longer peptides to facilitate direct comparison with the DnaJB4 results. One interesting finding in this study is that the P3 and P4 glutamates, which are strictly conserved in the EEVD motifs, are dispensable for binding to CHIP. In contrast, these two positions are involved in binding to DnaJB4 (see Fig 4C). In potential support of this idea, the P3 side chain of the EEVD is observed to make contacts with lysine 177 in the co-structure with DnaJB1’s CTD I. Thus, conservation of these glutamate residues might be primarily guided by the JDP interaction.

**Figure 5:**
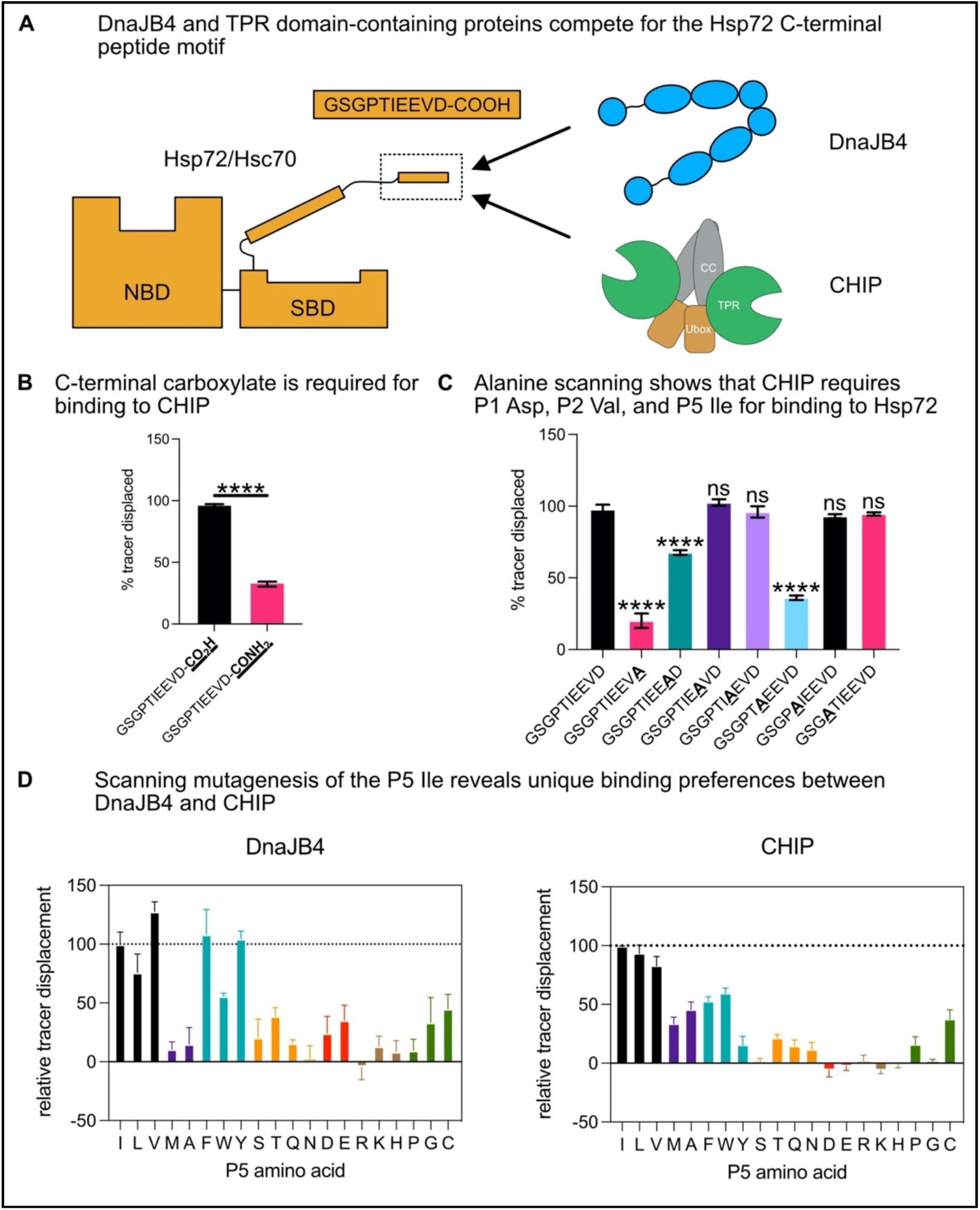
DnaJB4 accommodates an expanded number of amino acids at P5, compared to CHIP. (A) Cartoon depicting competition for the Hsp70 C-terminal EEVD motif by DnaJB4 and CHIP. (B) Competition FP experiment comparing displacement of 20 nM Hsp72 tracer from 1.58 µM CHIP by 100 µM WT or C-terminally amidated Hsp72 peptides. Graph shows mean tracer displacement relative to DMSO control ± SD (n=4). Statistics were performed using unpaired student’s t-test (****p<0.0001 compared to WT control). (C) Competition FP experiment comparing alanine scanning substitutions across the Hsp72 C-terminal sequence in binding to CHIP. Graph shows mean tracer displacement relative to DMSO control ± SD (n=4). Statistics were performed using unpaired student’s t-test (****p<0.0001, ns = not significant compared to WT control). (D). Competition FP experiment comparing all possible mutations at the P5 position of the Hsp72 EEVD motif in binding to DnaJB4 or CHIP. 5 µM DnaJB4 or 1.58 µM CHIP were incubated with 20 nM WT Hsp72 tracer and 100 µM unlabeled competitor peptide. Tracer displacement was calculated relative to DMSO control and normalized to WT as 100% displacement. Graph shows mean relative tracer displacement ± SD (n=4).

Next, to better understand the specific contributions at P5 for binding to both CHIP and DnaJB4, we tested Hsp72 peptides containing all natural amino acids at this position. With respect to CHIP binding, no substitution surpassed the native isoleucine in affinity. Further, only branched chain aliphatic residues (leucine and valine) were able to substitute at this position, while charged and polar residues were strongly disfavored (Fig 5D). With respect to DnaJB4, leucine and valine were able to substitute for isoleucine; further, valine substitution modestly enhanced binding. Like CHIP, charged and polar residues were largely disfavored at the P5 position. Strikingly, though, certain aromatic residues (phenylalanine and tyrosine) could substitute for isoleucine. As previously mentioned, the P5 isoleucine is thought to project into a hydrophobic pocket created by residues Ile231 and Phe233 of DnaJB4 (see Fig 2A). These residues are approximately 4 Å apart from one another, potentially allowing space for larger side chains. Thus, we hypothesize that phenylalanine or tyrosine may be accommodated and that they could potentially engage in pi-stacking interactions with Phe233 in this pocket. Conversely, the corresponding hydrophobic shelf in CHIP’s TPR domain is relatively narrow, which likely limits binding to small, branched-chain aliphatic residues^36^.

Collectively, these experiments suggest that DnaJB4 and CHIP use partially over-lapping molecular features to bind the EEVD motif. Specifically, DnaJB4 primarily recognizes the carboxylate and the P5 position, with an expanded preference for either branched aliphatic or small aromatic side chains. CHIP also recognizes the carboxylate and the P5 position, but it also makes key contacts with the P1/P2 residues and has a narrower requirement at P5.

### Mutations in the EEVD motif reduce collaboration between Hsp72 and DnaJB4

Pioneering work by the Craig group showed that the interaction of Hsp70’s EEVD with Class B JDPs is important for chaperone function^53,54^ and more recent structural studies have revealed that this effect is mediated by an allosteric release of autoinhibition that promotes JD function^39,55^. Here, we wanted to leverage our knowledge of DnaJB4’s SAR to probe these functional relationships in more detail. Towards this goal, we generated a mutant of full length Hsp72 in which the EEVD motif was deleted (Hsp72 ΔEEVD). Additionally, we created a point mutation of the critical P5 isoleucine residue (Hsp72 I637A), which significantly weakened, but did not abolish, binding to DnaJB4 (see Fig 4). These two mutants were then tested for their ability to collaborate with DnaJB4 in ATPase and luciferase refolding assays. First, we measured the intrinsic ATPase activities of the Hsp72 variants to create a baseline. As shown previously^21,53^, Hsp72 ΔEEVD had reduced intrinsic ATPase activity compared to the WT (WT = 10.9 ± 2.8 pmol ATP/min; ΔEEVD = 4.6 ± 1.9 pmol ATP/min). However, Hsp72 I637A had normal intrinsic activity (I637A = 15.8 ± 3.9 pmol ATP/min), so this mutant seemed better positioned for isolating the impact of DnaJB4 binding without the complicating effects on intrinsic turnover. Accordingly, we then measured the ability of DnaJB4 to stimulate ATPase activity by the Hsp72 variants. As expected, DnaJB4 stimulated the maximum ATPase activity (V_max,app_) of WT Hsp72 by ∼4-fold (37.8 ± 3.5 pmol ATP/min), at a half-maximal concentration (Km_app_) of ∼ 0.06 µM. Conversely, DnaJB4 was unable to stimulate the Hsp72 ΔEEVD mutant (Fig 6A), confirming previous reports that used other Class B JDPs^53^. Hsp72 I637A showed an intermediate level of activation, with a V_max,app_ of only ∼2-fold above baseline (23.9 ± 3.6 pmol ATP/min) and K_m,app_, ∼ 0.01 µM. Together, these results show that the affinity of the EEVD-CTD I interaction is important for ATP turnover.

**Figure 6:**
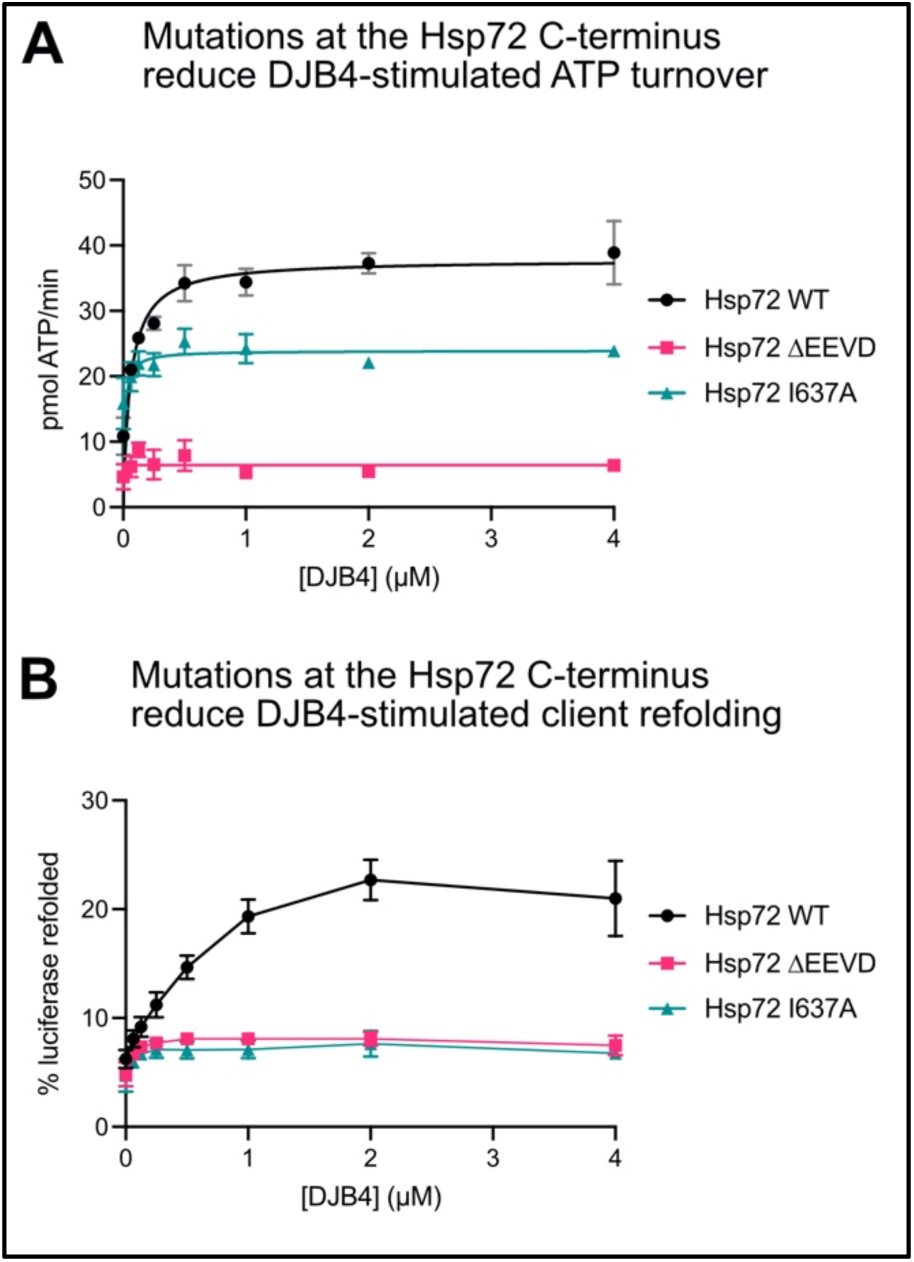
Mutations in the EEVD motif reduce collaboration between Hsp72 and DnaJB4. (A) ATP hydrolysis assay comparing the turnover rate of Hsp72 WT and mutants in the presence of DnaJB4, as measured by malachite green. The graph shows mean ATPase rate ± SD (n=3). Curves were fit according to Michaelis-Menten kinetics at steady-state. (B) Luciferase refolding assay comparing WT and mutant Hsp72 in the presence of DnaJB4. Refolding was measured by SteadyGlo luciferase reagent (see the methods for details). The graph shows mean percent luciferase refolded relative to non-denatured luciferase control ± SD (n=3).

Interestingly, the effects of the I637A mutant were even more pronounced in the luciferase refolding assay (Fig 6B), in which both Hsp72 ΔEEVD and I637A were nearly completely impaired in the ability to coordinate with DnaJB4. Thus, the EEVD interaction is absolutely required to promote client refolding, such that even the single isoleucine to alanine mutant could completely abrogate it. We speculate that this activity requires finely tuned kinetics. For example, DnaJB4 residence times may be shorter on the AEEVD motif compared to the IEEVD motif, leading to lower probability of proper coordination with Hsp72 during engagements with denatured luciferase.

### Competition for the EEVD motif by co-chaperones regulates chaperone functions

The complex of Hsp70 with CHIP is known to mediate the ubiquitination and degradation of client proteins^18,25^. Conversely, the complex of Hsp70 and DnaJB4 is most often associated with pro-folding functions^8^. Thus, we hypothesized that competition between the co-chaperones might reciprocally inhibit these distinct functions. Moreover, our studies had shown that the affinities of the EEVD motif for CHIP and DnaJB4 are similar *in vitro* (CHIP Kd ∼ 1 µM, DnaJB4 Kd ∼ 2 µM), suggesting that they might inhibit each other at near equivalent potency. To test this idea, we first performed ubiquitination assays, wherein CHIP was used to ubiquitinate Hsc70 or Hsp90α *in vitro*. Consistent with previous findings^60,61^, both Hsc70 and Hsp90 were robustly ubiquitinated by CHIP, creating the expected laddering of high molecular weight, ubiquitinated species (Fig 7A). Adding DnaJB4 to these mixtures resulted in dose-dependent inhibition of Hsc70 ubiquitination. As expected, DnaJB4 had no effect on Hsp90α ubiquitination, thus providing an important control.

**Figure 7:**
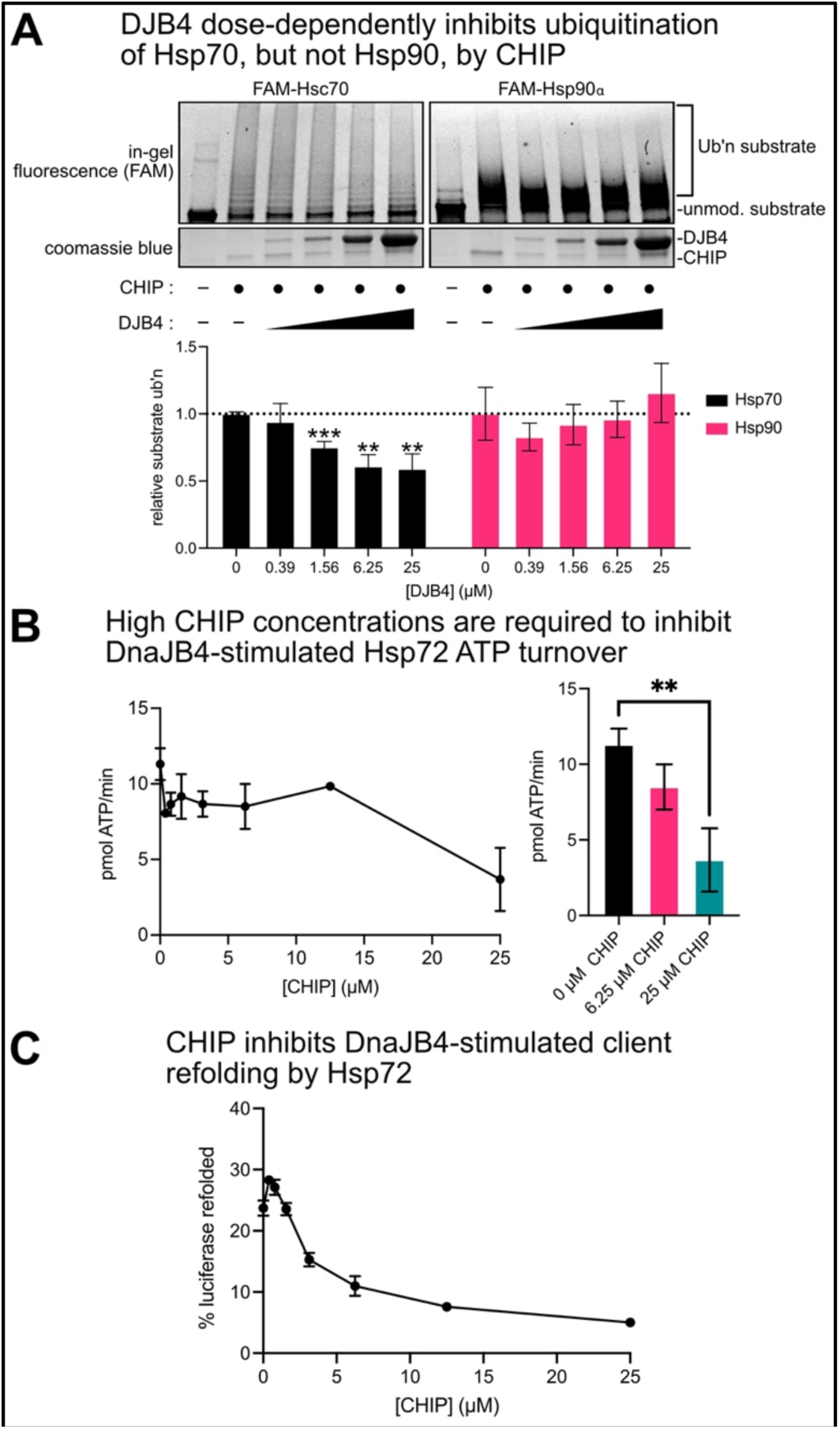
Competition for the EEVD motif by co-chaperones regulates chaperone functions. (A) In vitro ubiquitination assay comparing ubiquitination of Hsc70 or Hsp90α by CHIP in the presence of increasing amounts of DnaJB4. CHIP, FAM-labeled chaperone substrate, and increasing concentrations of DnaJB4 were incubated together for 30 min at room temperature prior to addition of ubiquitination machinery (Ube1, UbcH5, ubiquitin) and ATP. Reactions were performed for 10 min at room temperature, quenched in SDS-PAGE loading buffer, and denatured at 95 °C. Samples were separated by SDS-PAGE and ubiquitination was analyzed by in-gel fluorescence, while CHIP and DnaJB4 were identified by staining with Coomassie blue. The graph shows mean substrate ubiquitination relative to no DnaJB4 control ± SD (n=3). Statistics were performed using unpaired student’s t-test (**p<0.01, ***p<0.001 compared to no DnaJB4 control). (B) ATP hydrolysis assay comparing ATP turnover rate of WT Hsp72 in the presence of constant DnaJB4 and increasing concentrations of CHIP. Hsp72, DnaJB4, and increasing concentrations of CHIP were incubated with ATP for 1 hour at 37 °C and ATP hydrolysis was assayed using malachite green reagent. Graph shows mean ATP hydrolysis rate ± sd (n=3). Statistical analysis at right was performed using unpaired student’s t-test (**p<0.01). (C) Client refolding assay comparing ability of Hsp72 to refold client in the presence of DnaJB4 and CHIP. Hsp72 was incubated with denatured luciferase, DnaJB4, and increasing concentrations of CHIP for 1 hour at 37 °C, and client refolding was assayed by addition of SteadyGlo luciferase reagent. The graph shows mean percent luciferase refolded relative to non-denatured luciferase control ± SD (n=3).

Next, we explored whether CHIP might interrupt DnaJB4’s ability to stimulate Hsp72’s ATPase activity. We found that 40-fold excess of CHIP (25 µM) relative to DnaJB4 (625 nM) was required to observed significant inhibition (Fig 7B). This finding matches with previous observations, in which excess CHIP was required to block the activity of DnaJB1 in similar assays^62^. Relatively high concentrations of CHIP may be required to suppress DnaJB4 function because multivalent contacts between Hsp70 and DnaJB4, mediated by both the JD and CTD I, effectively enhance avidity.

We subsequently tested the ability of CHIP to suppress client refolding by the Hsp72-DnaJB4 complex. Indeed, titrations of CHIP into folding reactions showed that it is a potent inhibitor (Fig 7C). The more pronounced ability of CHIP to suppress client refolding, compared to ATPase activity, is likely influenced by several factors, including CHIP’s described function as a “holdase” that can bind directly to unfolded clients, as well as CHIP’s preference for Hsp72 in the closed, ADP-bound state^63,64^. Together, these studies suggest that CHIP and DnaJB4 compete for the EEVD motif to tune formation of Hsp70 complexes and influence chaperone function.

### Pseudo-phosphorylation of the Hsp70 C-terminus inhibits CHIP binding but has no effect on DnaJB4

Molecular chaperones are subject to myriad post-translational modifications (PTMs), including AMPylation, methylation, acetylation, phosphorylation, and others^65,66^. Moreover, some PTMs have been directly linked to changes in the binding to co-chaperones. Specifically, phosphorylation of the P6 threonine residue near the Hsp70 EEVD motif is known to inhibit binding of CHIP^30,67^. We confirmed this effect in our hands, as the mutation of P6 to the phosphomimetic, glutamic acid, resulted in a 5-fold weakening of the affinity of an Hsp72 peptide for CHIP (Fig 8A). In contrast, the same peptide had no effect on binding to DnaJB4 (Fig 8A), suggesting that phosphorylation might have a selective effect on CHIP but not DnaJB4. This observation is also supported by examination of the predicted binding modes for the EEVD motif when bound to CHIP or DnaJB4 (Fig 8B, PDB 6EFK^36^ and 3AGY^37^). When bound to CHIP, the EEVD motif is configured into an unstructured, bent conformation and it engages in multiple sidechain interactions via the TPR domain^27^. Notably, the P6 threonine is engaged in a hydrogen bonding interaction with the TPR domain, and phosphorylation of this residue is likely to generate electrostatic and/or steric clashes^36^. Conversely, when bound to DnaJB4, the EEVD motif adopts a beta sheet conformation with the P6 threonine being relatively exposed to solvent^37^. Phosphorylation of this residue is therefore less likely to modulate binding to DnaJB4’s CTD I, consistent with the FP studies. Together, these results suggest that cells could use PTMs, especially phosphorylation of the C-terminus of Hsp70s, to further tune binding at this PPI hotspot. More broadly, the drastically different configurations of the EEVD motif (*e*.*g*., “bent” vs. “linear”) when bound to these two domains further highlights the idea that molecular recognition by CHIP and DnaJB4 relies on only partially over-lapping molecular features.

**Figure 8:**
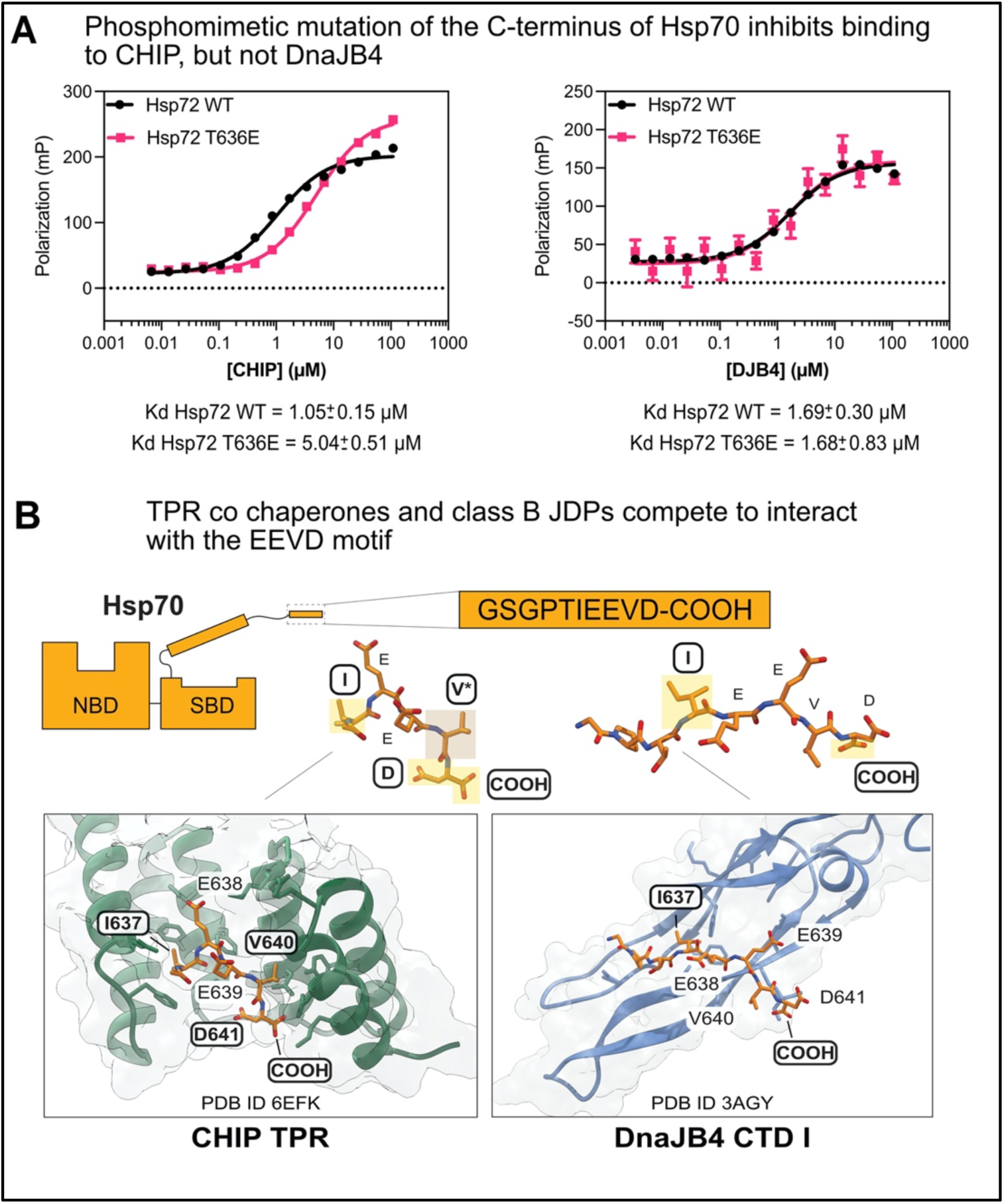
Pseudo-phosphorylation of the Hsp70 C-terminus inhibits CHIP binding but has no effect on DnaJB4. (A) Saturation binding FP experiments comparing WT or phosphomimetic T636E Hsp72 fluorescent tracer binding to CHIP (left) or DnaJB4 (right). Increasing concentrations of CHIP or DnaJB4 were incubated with Hsp72 fluorescent tracer for 30 minutes at room temp. The results are the average of four replicates and error bars represent SD. The Kd values for each condition are expressed as mean with a 95% CI. (B) Comparison between the binding modes of Hsp70’s EEVD motif interacting with CHIP and DnaJB4. The CHIP TPR (green) orients the Hsp70 EEVD motif in a hooked or bent conformation, where it makes interactions with the P5 Ile, P2 Val, P1 Asp, and C-terminal carboxylate. Conversely, CTD I of DnaJB4 (blue) binds the Hsp70 EEVD motif in a linear orientation, with the P5 Ile and C-terminal carboxylate make important contacts. Residues important for binding to each co-chaperone are highlighted on the crystal structures (CHIP TPR-Hsp70 EEVD: PDB 6EFK^36^, DnaJB1 CTD I-Hsp70 EEVD: PDB 3AGY^37^).

## Discussion

Interactions of Hsp70 with its co-chaperones impart a strikingly diverse set of cellular functions to this molecular chaperone, allowing it to act in client folding, trafficking and degradation. Thus, a major goal in the proteostasis field is to understand when and where a particular complex between Hsp70 and its co-chaperones will assemble. This is a challenging problem because there are approximately 13 NEFs^9^, 44 JDPs^8^, and 35 TPR co-chaperone genes^28^ and when these factors are combined with the 6 cytosolic Hsp70s^68^, an upper limit of >120,000 unique possible combinations are possible. While the true number of complexes is likely much lower than this value because of restrictions in subcellular localization and tissue-specific expression, studies have supported the broad idea that cells contain many Hsp70 complexes^69–71^. Thus, it is important to understand which co-chaperones might compete and which molecular determinants are used to drive these decisions.

Here, we focused on studying how the TPR domain protein, CHIP, and Class B JDP, DnaJB4, converge on Hsp72’s EEVD motif (see Figs 2B and 5A). This set of PPIs seemed especially important to understand because these co-chaperones promote opposing functions of Hsp70, with DnaJB4 directing the client to a pro-folding pathway and CHIP favoring client destruction^53,62^. Thus, competition for binding the EEVD could be central to the triage decisions made by the Hsp70 system. Indeed, we observed reciprocal inhibition of Hsp70’s functions (see Figs 7A and 7B), suggesting that distinct classes of co-chaperones regulate the functional outcomes of others via competition for the EEVD motif.

What controls the “decision” of Hsp70 to bind CHIP vs DnaJB4? It is easy to imagine that (at least) two parameters – relative affinity for the EEVD motif and relative abundance of a particular co-chaperone – would combine to dictate which partner would bind at this PPI “hotspot”. Under the conditions tested, we found that CHIP has a slightly tighter affinity for the EEVD motif than DnaJB4 (see Fig 2A, Fig 8A). Moreover, CHIP is abundant, constitutively expressed, and has minimal tissue specificity^17,72^. These observations would suggest that CHIP is typically more available for binding to the EEVD motif, thereby potentially favoring client clearance over client folding. DnaJB4, however, is inducible under proteotoxic stress, greatly boosting its expression^73,74^. Additionally, we observed a significant weakening of CHIP’s affinity when a phosphomimetic mutation is added to the EEVD motif, while DnaJB4 was unaffected (see Fig 8A). Thus, signal transduction via transcription or phosphorylation would seem likely to favor DnaJB4 binding over CHIP. This could be why previous studies have observed that CHIP over-expression does not lead to client degradation, as might otherwise be predicted^75^. Finally, we also observed tighter binding of DnaJB4 to the constitutive Hsc70 versus the stress-inducible Hsp72 (see Fig 3B, 3C), and a dose-dependent protection of Hsc70 from CHIP-dependent ubiquitination by DnaJB4 (see Fig 7A). Thus, the relative levels of Hsc70 and Hsp72 might also dictate which complexes are formed and how quickly the chaperones are turned over. Finally, the relative kinetics of the EEVD interactions with TPR proteins and JDPs are not yet clear. Because Hsp70’s functions require careful coordination of multiple, weak binding events^76^, the relative association/dissociation rates and co-chaperone residence times are likely to be important parameters, dictating both which complexes are formed and what allosteric signals are transmitted through those complexes. This may explain why ATPase stimulation of the Hsp72 I637A mutant is hampered; changes in binding kinetics due to the mutation may lead to a lower probability of allosteric signal transduction to the distal Hsp72 NBD. Together, these findings suggest how cells might employ PTMs and transcriptional responses to fine-tune co-chaperone affinities and concentrations, dictating which complexes are favored and, in turn, what Hsp70 functions are favored. In addition, this discussion must also clearly acknowledge that there are many other TPR and JDP co-chaperones in cells (besides CHIP and DnaJB4), which provide additional layers of competition for the EEVD motifs.

Certain client proteins are also likely to tune these PPIs. For example, clients have been shown to bind CTD I and CTD II of JDPs^46,49^, such that they would be expected to potentially compete with Hsp70’s EEVD motif. Accordingly, the production of unfolded clients by proteotoxic stress may directly impede EEVD-binding to Class B JDPs, perhaps promoting the formation of Hsp70-CHIP complexes. CHIP, on the other hand, directly interacts with a subset of substrates that are generated by caspase-dependent proteolysis^36^. Briefly, caspase activity produces new C-termini that end in an aspartic acid, and some of these can resemble the EEVD motif. While EEVD binding to CHIP requires a C-terminal aspartate (see Fig 5C), however, we found that EEVD binding to DnaJB4 does not require this sidechain (see Fig 4C), suggesting that CHIP is more selective for neo-C-termini generated by caspase cleavage. Therefore, caspase activation may selectively displace CHIP, but not DnaJB4, from Hsp70. These scenarios highlight likely roles for clients in further shaping the distribution of proteostasis complexes in cells.

Chemical probes that can selectively perturb chaperone -co-chaperone PPIs are desirable tools for dissecting the role of these complexes in cellular functions^77^. Effective probes of this type would benefit from the ability to differentiate between closely related PPIs, such as those between the EEVD motif and either CHIP or DnaJB4. Thus, we were interested in the finding that the EEVD motif binds to CHIP and DnaJB4 with partially distinct structural features. Specifically, the expanded side chain preferences of DnaJB4 for the P5 residue and its reliance on the P3 and P4 glutamates suggest that small molecules might be able to preferentially block EEVD binding to this co-chaperone over others. On the other hand, the requirement for a P1 aspartic acid and P2 valine in binding to CHIP, but not for DnaJB4, likewise presents a potential opportunity for selectivity (see Fig 4C). These predictions will require additional exploration, but it is compelling that the two classes of co-chaperones “read” partially different chemical information in the EEVD motif.

## Methods

### Plasmids

All recombinant proteins were expressed from a pMCSG7 vector with an N-terminal 6-His tag and TEV protease cleavage site.

### Protein expression and purification

#### DnaJB4

DnaJB4 was expressed in *E. coli* BL21 (DE3) Rosetta (New England BioLabs) cells. Liter cultures of terrific broth were grown at 37 °C until the OD_600_ reached 0.8. Cultures were then cooled to 18 °C, induced with 500 µM isopropyl beta-D-1-thiogalactopyranoside (IPTG) and grown overnight at 18 °C. Cell pellets were resuspended in His binding buffer (50 mM Tris pH 8.0, 10 mM imidazole, 750 mM NaCl) supplemented with cOmplete EDTA-free protease inhibitor cocktail (Sigma-Aldrich). Cells were lysed by sonication and pelleted by centrifugation, and the supernatant was applied to a 5 mL HisTrap Ni-NTA Crude column (Thermo Fisher Scientific). The column was washed with His binding buffer, followed by His wash buffer 1 (50 mM Tris pH 8.0, 30 mM imidazole, 750 mM NaCl, 3% EtOH) and His wash buffer 2 (50 mM Tris pH 8.0, 30 mM imidazole, 100 mM NaCl, 3% EtOH) supplemented with 1 mM ADP. The protein was eluted with a gradient elution from 0-100% His elution buffer (50 mM Tris pH 8.0, 300 mM imidazole, 300 mM NaCl). Eluent was supplemented with 1 mM DTT and TEV protease to remove the N-terminal His tag, and cleavage was allowed to proceed overnight at 4 °C and dialyzed to His Binding buffer. The protein was then buffer exchanged into His binding buffer and applied to Ni-NTA His-Bind Resin to remove His-tagged TEV protease. The protein was further purified by size exclusion chromatography (SEC) using an AKTA Pure chromatography instrument (Cytiva) using Superdex 200 column (Cytiva) in Tris buffer (50 mM Tris pH 8.0, 300 mM NaCl).

#### Hsp70s

WT and mutant Hsp72/HSPA1A and Hsc70/HSPA8 were expressed in *E. coli* BL21(DE3) Rosetta cells (or BL21(DE3) for point mutants). Liter cultures of terrific broth (TB) were grown at 37 °C until an OD_600_ value of 0.6. Cultures were cooled to 20 °C and induced with 500 µM IPTG. Cultures were then grown overnight at 20°C. Cell pellets were resuspended in binding buffer (50 mM Tris pH 8.0, 10 mM imidazole, 500 mM NaCl) supplemented with cOmplete EDTA-free protease inhibitor cocktail (Sigma-Aldrich). Cells were lysed by sonication and pelleted by centrifugation, and the supernatant was applied to HisPur Ni-NTA resin (Thermo Fisher Scientific). The resin was washed with binding buffer, washing buffer (50 mM Tris pH 8.0, 30 mM imidazole, 300 mM NaCl), and protein was eluted with elution buffer (50 mM Tris pH 8.0, 300 mM imidazole, 300 mM NaCl). Eluent was supplemented with 1 mM DTT and TEV protease to remove the N-terminal His tag, and cleavage was allowed to proceed overnight at 4 °C. The protein was applied to column packed with ATP-agarose (Sigma Aldrich) and the column was washed with buffer A (25 mM HEPES pH 7.5, 5 mM MgCl_2_, 10mM KCl) and buffer B (25 mM HEPES pH 7.5, 5 mM MgCl_2_, 1M KCl). Protein was eluted with buffer A supplemented with 3 mM ATP.

#### CHIP

Recombinant human CHIP was expressed in BL21(DE3) (New England Biolabs) *E. coli* and grown in terrific broth (TB) to OD_600_ = 0.6 at 37 °C. Cells were cooled to 18 °C, induced with 500 µM isopropyl β-D-1-thiogalactopyranoside (IPTG), and grown overnight. Cells were collected by centrifugation, resuspended in binding buffer (50 mM Tris pH 8.0, 10 mM imidazole, 500 mM NaCl) supplemented with protease inhibitors, and sonicated. The resulting lysate was clarified by centrifugation and the supernatant was applied to Ni^2+^ -NTA His-Bind Resin (Novagen). Resin was washed with binding buffer and His wash buffer (50 mM Tris pH 8.0, 30 mM imidazole, 300 mM NaCl), and then eluted from the resin in His elution buffer (50 mM Tris pH 8.0, 300 mM imidazole, 300 mM NaCl). Following, the N-terminal His tag was cleaved by overnight dialysis with TEV protease at 4 °C. Digested material was applied to His-Bind resin to remove cleaved His tag, undigested material and TEV protease. Protein was further purified by SEC in CHIP storage buffer (50 mM HEPES pH 7.4, 10 mM NaCl), concentrated, flash frozen in liquid nitrogen, and stored at -80 °C.

### Peptides

Peptides were ordered from GenScript (95% purity by high performance liquid chromatography (HPLC)). Fluorescence polarization tracer peptides were designed with a 5-carboxyfluorescein (5-FAM) moiety linked to the peptide N-terminus via a six-carbon spacer (aminohexanoic acid). Unlabeled peptides were N-terminally acetylated. Unless specified, peptides bore an unmodified free carboxylate at the C-terminus. Peptides were diluted in DMSO to 10 mM stock solutions and stored at -20 °C.

### Fluorescence polarization

#### General

All FP experiments were performed in 384-well, black, low-volume, round-bottom plates at a final assay volume of 18 µL (Corning 4511). Polarization values in millipolarization units (mP) were measured at an excitation wavelength of 485 nm and an emission wavelength of 525 nm, with 100 flashes per read using a Spectramax M5 plate reader (Molecular Devices). All experiments were performed 2 times in quadruplicate. Experimental data were analyzed using GraphPad Prism 9. Saturation binding data was background subtracted, and curves were fit using the model [Agonist] vs. response (three parameters). For competition experiments, data was background subtracted to tracer alone and normalized to DMSO control to determine relative tracer displacement.

#### Saturation binding to Hsp72 C-terminal probes

A sample of 20 nM Hsp72 tracers (FITC-Ahx-GSGPTIEEVD or FITC-Ahx-GSGPEIEEVD) was incubated with various concentrations of DnaJB4 (WT or QPD mutant) or CHIP in 2X dilutions (final [protein] = 110 to 0.013 µM) in binding buffer (JDP binding buffer = 50 mM Tris, 15 mM NaCl, 1 mM DTT, 0.01% Triton X-100, pH 8.0; CHIP binding buffer = 50 mM HEPES, 50 mM KCl, 0.01% Triton X-100, pH 7.4). The plate was covered from light and allowed to incubate at room temperature for 30 min prior to reading.

#### DnaJB4 FP competition experiments

Unlabeled peptides were assessed for the ability to compete with the Hsp72 tracer. Briefly, 100 µM peptides were incubated with 5 µM DnaJB4 and 20 nM Hsp72 tracer in JDP binding buffer (see above). The plate was covered from light and allowed to incubate at room temperature for 30 min prior to reading.

#### CHIP FP competition experiments

Mixtures of 1.58 µM CHIP and 20 nM Hsp72 tracer were incubated with 100 µM unlabeled competitor peptides in CHIP FP assay buffer (50 mM HEPES pH 7.4, 50 mM KCl, 0.01% Triton X-100). The plate was covered from light and allowed to incubate at room temperature for 30 min prior to reading.

### Differential scanning fluorimetry

DSF was performed with a 15 µL assay volume in 384-well Axygen quantitative PCR plates (Fisher Scientific) on a qTower^3^ real-time PCR thermal cycler (Analytik Jena). Fluorescence intensity readings were taken over 70 cycles in “up-down” mode, where reactions were heated to desired temp and then cooled to 25 °C before reading. Temperature was increased 1 °C per cycle. Each well contained 5 µM DnaJB4, 5× Sypro Orange dye (Thermo Fisher), and 100 µM of peptide in JDP binding buffer. Fluorescence intensity data was truncated between 45 -70 °C, plotted relative to temperature, and fit to a Boltzmann Sigmoid in Prism 9.0 (GraphPad). DnaJB4 apparent melting temp (Tm_app_) was calculated based on the following equation: Y=Bottom+((Top-Bottom)/(1+exp(*Tm*-*T*/Slope)))

### ATP Hydrolysis Assays

ATPase assays were carried out using the malachite green assay as described^14,78^. In brief, 1 *µ*M Hsp72 and various concentrations of DnaJB4 were added to clear 96-well plates, and the reactions were initiated by addition of 2.5 mM ATP. Reactions were allowed to proceed for 1 hour at 37 °C, after which they were developed using malachite green reagent and quenched with sodium citrate. Plate absorbance was measured at 620 nm, and a standard curve of sodium phosphate was used to convert the absorbance values to pmol ATP/μM Hsp72/min. V_max_ and K_m, app_ were derived as fit parameters to a modified Michaelis-Menten model (ATPase rate=V_max_ *[DnaJB4]/(K_m, app_ +[DnaJB4])) where V_max_ reflects the maximal increased ATP hydrolysis conferred by DnaJB4 binding and K_m,app_ represents the half-maximal concentration of DnaJB4 binding to and stimulating the ATPase activity of Hsp72.

### Luciferase Refolding Assays

Experiments were performed as described^14^. Briefly, *Renilla* luciferase (Promega) was denatured in 6M GdnHCl for 1 hour at room temperature. Hsp72 and denatured luciferase were diluted into a working concentration in buffer containing an ATP regenerating system (23 mM HEPES, 120 mM KAc, 1.2 mM MgAc, 15 mM DTT, 61 mM creatine phosphate, 35 units/mL creatine kinase, and 5 ng/µL BSA, pH 7.4). A titration series of DnaJB4 was added, and the reaction was initiated with the addition of 2.5 mM ATP. The assay proceeded for 1 hour at 37 °C in white, 96-well plates, and luminescence was measured using the SteadyGlo luminescence reagent (Promega).

### *In vitro* ubiquitination assays

Four 4× stock solutions were prepared containing (i) Ube1 (R&D Systems) + UbcH5c/UBE2D3 (R&D Systems) (400 nM Ube1 and 4 μM UbcH5c), (ii) Ubiquitin (R&D Systems) (1 mM Ub), (iii) CHIP + substrate + DnaJB4 (4 μM CHIP, 4 μM substrate, varying concentrations of DnaJB4) and (iv) ATP + MgCl_2_ (10 mM ATP and 10 mM MgCl_2_) in ubiquitination assay buffer (50 mM Tris pH 8.0, 15 mM NaCl). CHIP + substrate + DnaJB4 solutions were allowed to equilibrate at room temperature for 30 min prior to initiating the reaction. Ubiquitination reactions were generated by adding 10 μL of each 4× stock, in order from 1 to 4, for a final volume of 40 μL(100 nM Ube1, 1 μM UbcH5c, 250 μM ubiquitin, 2.5 mM ATP, 2.5 mM MgCl_2_, 1 μM CHIP and 1 μM substrate). Reactions were incubated at room temperature for 10 min, quenched in 20 µL 3× SDS–PAGE loading buffer (188 mM Tris-HCl pH 6.8, 3% SDS, 30% glycerol, 0.01% bromophenol blue, 15% β-mercaptoethanol), and heated to 95 °C for 5 min. Samples were separated by SDS-PAGE on 4-20% polyacrylamide gels (BioRad) and analyzed by in-gel fluorescence on a Chemidoc Imager (BioRad) using the SYBR green fluorescence setting or by staining with Coomassie blue reagent. Quantitation of substrate ubiquitination was performed by densitometry analysis in ImageJ (NIH), which was thresholded to a no-CHIP control.

### Protein Labeling with 6-Carboxyfluorescein

Substrates for *in vitro* ubiquitination assays were labeled with 6-carboxyfluorescein (FAM) to enable in-gel fluorescence measurement of ubiquitination as previously described^36^. Briefly, proteins were dialyzed into labeling buffer (25 mM HEPES pH 7.4, 50 mM KCl, 1 mM TCEP) and labeled by addition of 5 eq. of maleimide-FAM (Fisher Scientific 501143190) for 2 hours at room temperature. The reaction was quenched by addition of 1 mM DTT, and excess reagent was removed by iterative concentration and dilution over a 10 kDa MWCO microcentrifuge spin column (Pierce).

## Author Contributions

CMN generated reagents, designed and performed experiments, analyzed data, and drafted the manuscript. OTJ generated reagents, designed and performed experiments, and analyzed data. ECC generated reagents and performed experiments. TA generated reagents, designed and performed experiments. JEG designed experiments and drafted the manuscript. All authors edited the manuscript.

## Data Availability

All data are in the manuscript. Source data is available upon request from the corresponding authors: Taylor Arhar, or Jason E. Gestwicki.

## Acknowledgements

The authors acknowledge support from the NIH (OTJ K12GM081266; JEG NS059690 and AG068125). Recombinant Hsp90α was provided as a gift from the laboratory of Daniel Southworth (UCSF).

